# Comment on ‘Initiation of chromosome replication controls both division and replication cycles in *E. coli* through a double-adder mechanism’

**DOI:** 10.1101/2020.05.08.084376

**Authors:** Guillaume Le Treut, Fangwei Si, Dongyang Li, Suckjoon Jun

**Affiliations:** Department of Physics, University of California, San Diego, La Jolla, CA 92093, USA; Section of Molecular Biology, Division of Biology, University of California, San Diego, La Jolla, CA 92093, USA

## Abstract

Witz *et al*. recently performed single-cell mother machine experiments to track growth and the replication cycle in *E. coli*. They analyzed the correlation structure of selected parameters using both their data and published data, and concluded that *E. coli* cell-size control is implemented at replication initiation, which challenged the newly emerged division-centric mechanism of cell-size control in bacteria. We repeated Witz et al.’s analysis, and performed additional experiments and analytical calculations. These results explain Witz et al.’s observation and in fact support the division-centric model.

## Introduction

Cell size control is one of the long-standing problems in biology (Jun et al. 2018). The recent discovery of the adder principle has demonstrated that many evolutionary divergent organisms in both bacteria and eukaryotes share the same homeostasis strategy for size control despite their differences in molecular details and biological complexity (Campos et al. 2014; Taheri-Araghi et al. 2015; Jun and Taheri-Araghi 2015). This makes bacteria an attractive and tractable system to understand the mechanistic origin of cell-size control.

An outstanding issue in bacterial cell-size control is about what physiological parameters size control is imposed on. For example, the celebrated Helmstetter-Cooper model established three physiological parameters are necessary and sufficient to completely describe the progression of bacterial cell cycle and cell size (Cooper and Helmstetter 1968). Helmstetter and Cooper chose the growth rate (λ), cell size at replication initiation (S_i_), and the time elapsed during DNA replication and the subsequent cell division (C+D) (Figure 1A). Therefore, their model is often interpreted as replication initiation being a major implementation point of cell-cycle control. Formally, however, other sets of three independent parameters (Figure 1A) could constitute a cell-size control model.

**Figure 1:**
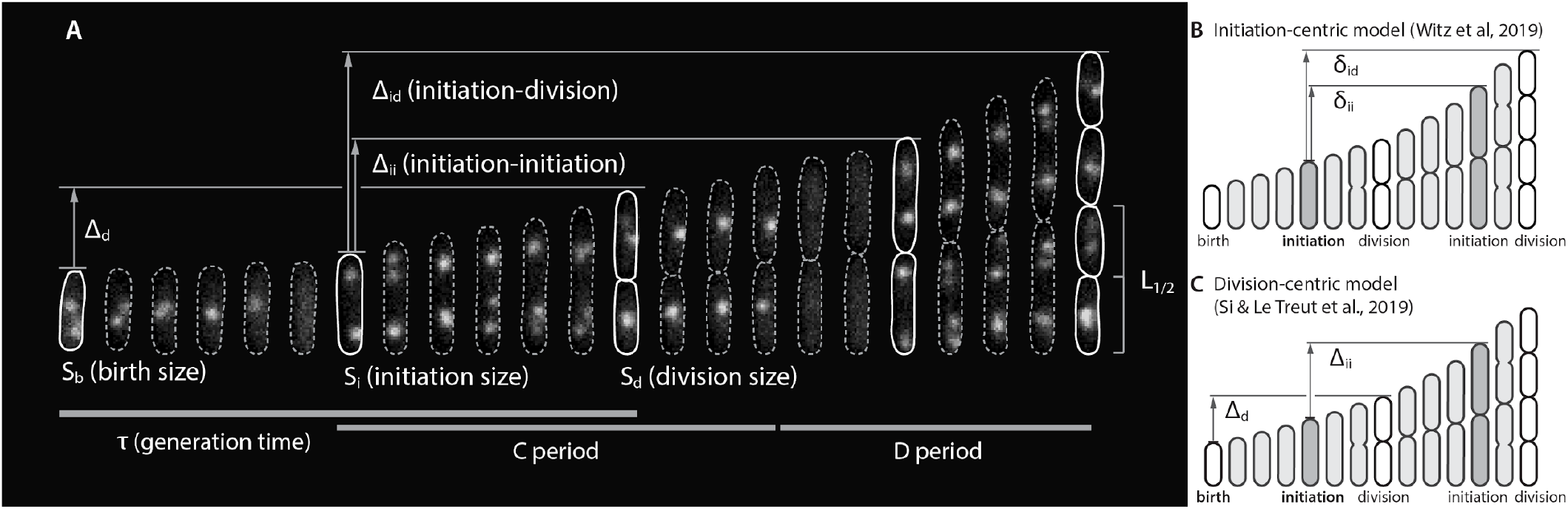
Physiological parameters that can be measured from single-cell experiments. **A**. The Helmstetter-Cooper model describes cell size and cell cycle using three parameters: generation time τ, C+D, and the initiation size S_i_. Additional parameters that can be measured include size added between two consecutive initiations Δ_ii_ = δ_ii_ x N_ori_, size added between two consecutive divisions Δ_d_, size added between initiation and division Δ_id_ = δ_id_ x N_ori_, relative septum position L_1/2_, and the elongation rate λ = dln(*l*)/d*t*, where *l* is the cell length (not shown). **B**. The initiation-centric model by Witz et al. (2019) proposed λ, δ_ii_, and δ_id_ (lower-cases δ’s indicate size normalized per origin) as the indepently physiological parameters, whereas **C**. the division-centric model by Si & Le Treut et al. (2019) has proposed λ, δ_ii_, and Δ_d_ as independent physiological parameters. Note that δ_id_ can span multiple generations, as the C+D period can in the Helmstetter-Cooper model. The number of origins in Witz et al.’s initiation-centric model is therefore determined by δ_id_/δ_ii_, similar to (C+D)/τ in the Helmstetter-Cooper model. By contrast, in the division-centric model by Si and Le Treut et al., the added sizes Δ_d_ and Δ_ii_ are between two consecutive cell cycles.

The recent eLife paper by Witz et al. (Witz, van Nimwegen, and Julou 2019) revisited the question of implementation point for cell size control. To this end, they tracked replication and division cycles at the singlecell level, using methods similar to previous works (Wallden et al. 2016; Si and Le Treut et al. 2019; Adiciptaningrum et al. 2015). They computed correlations between all pairs of measured physiological parameters, and identified a set of most mutually uncorrelated parameters. They then assumed that statistically uncorrelated physiological parameters must represent biologically independent controls. Such approaches previously facilitated the discovery of the adder principle and its formal description (Taheri-Araghi et al. 2015). Based on the correlation analysis, Witz et al. concluded that cell size at replication initiation is the most likely implementation point of size control, supporting previous theoretical models of initiation-centric size control (Amir 2014, 2017) but directly contradicting the division-centric mechanism revealed in our previous work (Si and Le Treut et al. 2019).

Here, we repeated Witz et al.’s analysis and also performed additional experiments and modeling as detailed below. We show that (1) Witz et al.’s own analysis of published data in fact supports the division-centric model. (2) Our new experiments using the *E. coli* BW strain that has the genetic background similar to Witz *et al*.’s also support the division-centric model. (3) In general, their correlation-based approach cannot distinguish different models. (4) Their initiation-centric double-adder model is analytically inconsistent with the adder phenotype and, in fact, produces a sizer-like behavior, making it an unlikely model of cell-size homeostasis in bacteria.

We expect the experiment, analysis, and modeling presented in this work to be a useful resource for researchers who are considering interdisciplinary, single-cell approaches to quantitative microbial physiology.

## Results

### Summary of Witz *et al*.’s correlation approach and results

We first summarize the results of correlation analysis in Witz et al. (2019). They considered a total of 10 measured parameters and 10 derived parameters, some of which are shown in Figure 1A. Since the work by Helmstetter and Cooper (1968), it has been known that the progression of cell size and cell cycle can be completely described using 3 variables (Wallden et al. 2016; Si et al. 2017). Therefore, there are at most 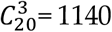 possible combinations. To measure the statistical independence of each set of parameters, Witz et al. constructed a correlation matrix (Figure 2).

**Figure 2:**
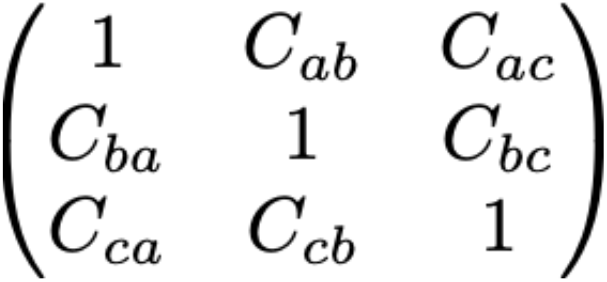
Correlation matrix.

The diagonal elements are 1’s and the off-diagonal elements are cross-correlations between pairs of parameters (a, b and c). Therefore, if all parameters are statistically independent of each other, the off-diagonal elements should be 0, and the determinant *I* of the matrix should be 1 (Figure 2). Based on this observation, Witz et al. used the determinant *I* ≤ 1 of the matrix as a metric for statistical independence of the hypothetical control parameters, with *I* = 1 being the set of most independent parameters. Note that Witz *et al*. analyzed a set of 4 parameters, instead of 3, out of the 20 aforementioned parameters to compute *I* (see next section and Materials and Methods).

To generate single-cell data, Witz *et al*. performed four mother machine experiments for three different growth conditions (Witz, van Nimwegen, and Julou 2019). In addition, they also chose to analyze one of the 9 nutrient-limitation experiment data sets from Si & Le Treut *et al*. (2019). They found the following four parameters to be statistically most independent and produce the highest value of *I*: λ (elongation rate), δ_ii_ (size added from initiation to initiation per origin), δ_id_ (size added from initiation to division per origin), and Λ_i_ (initiation mass per origin).

They concluded that replication initiation is the most upstream control of cell size (via δ_ii_ and Λ_i_), as division is triggered after growing by a constant size per origin since initiation (δ_id_) via an unknown mechanism (Figure 1B). This last point is initiation-centric and directly contradicts our previous findings that (1) replication and division are independently controlled via three parameters λ, δ_ii_ and Δ_d_ at the mechanistic level in both *Escherichia coli* and *Bacillus subtilis*; (2) it is cell division, not replication initiation, that drives cell-size homeostasis (Si and Le Treut et al. 2019).

### The result of Witz *et al*.’s correlation approach is more consistent with the division-centric model, rather than the initiation-centric model

Since the initiation-centric model and division-centric model are mutually exclusive (see next section for correlation analysis), we were initially puzzled that the experiment in Si & Le Treut et al. (2019) supported the initiation-centric model. However, this particular experiment was a special case, because that was the slow growth condition wherein *E. coli* deviated from the adder phenotype (Wallden et al. 2016; Si and Le Treut et al. 2019). This deviation is caused by active degradation of FtsZ by ClpXP in slow growth conditions (Männik, Walker, and Männik 2018; Sekar et al. 2018; Si and Le Treut et al. 2019), which we mechanistically explained by demonstrating that repression of clpX restores the adder phenotype (Si and Le Treut et al. 2019).

We therefore set out to re-analyze all our 9 steady-state nutrient-limitation growth experiments in *E. coli* and *B. subtilis* using the same approach Witz et al. used. We analyzed a set of 3 parameters as defined in Figure 1B and 1C, instead of 4 parameters. This difference in the number of parameters does not affect the overall conclusion of the *I*-value analysis (see Materials and Methods). We computed *I* values for three models: the “initiation-centric” adder model (Witz, van Nimwegen, and Julou 2019), the “division-centric” adder model (Si and Le Treut et al. 2019), and also the classic Helmstetter-Cooper model (Cooper and Helmstetter 1968) as a reference.

The results of the analysis are clear (Figure 3). Out of the total of nine experiments that we analyzed using Witz *et al*.’s method, six experiments were more consistent with the division-centric model (Si and Le Treut et al. 2019). On the other hand, both models performed consistently better than the Helmstetter-Cooper model.

**Figure 3:**
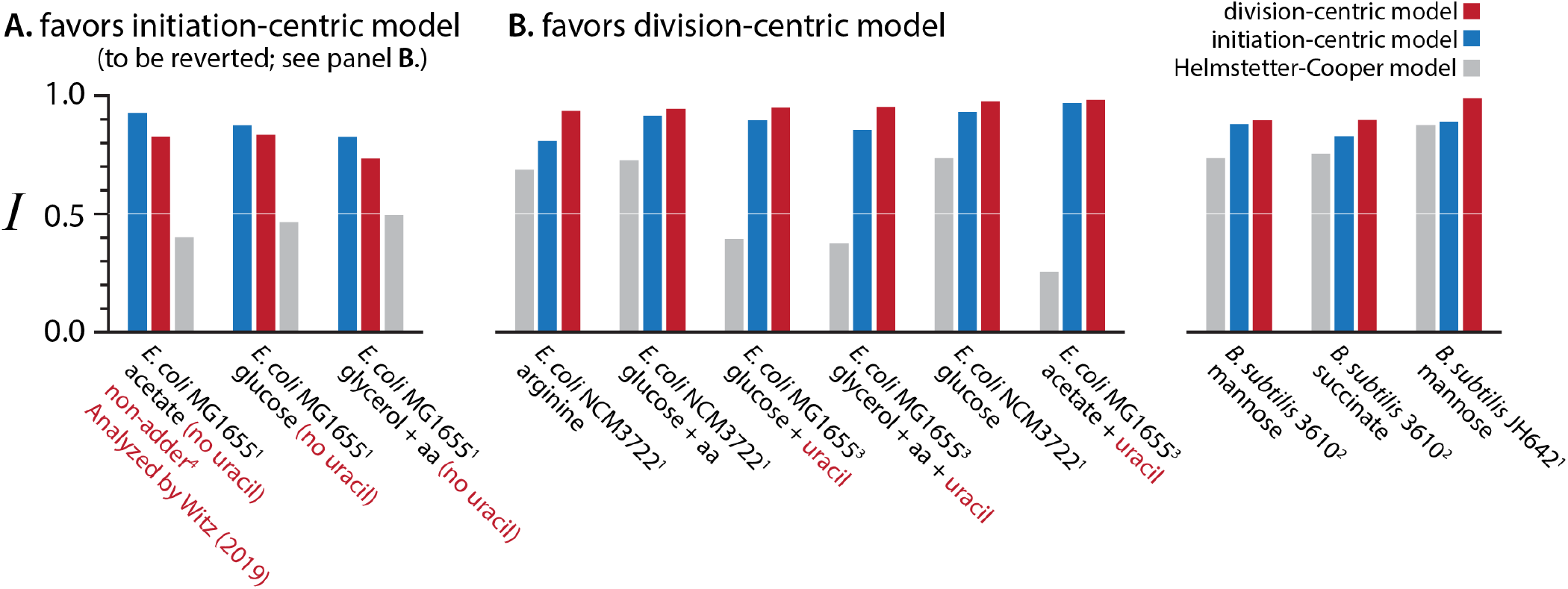
Data in *E. coli* and *B. subtilis* support the division-centric model (Si & Le Treut et al., 2019), rather than the initiation-centric model (Witz et al., 2019) or the Helmstetter-Cooper model (1968). (**A**). Our MG1655 data initially supported the initiation-centric model. However, this was due to the well-known pyrimidine pseudo-auxotrophy in MG1655 (Soupene et al. 2003). To alleviate the growth defected, we supplemented growth media with uracil, and the results reverted (**B**). All other published data supported the division-centric model. Superscripts represent the data sources: 1 - Si & Le Treut et al., 2019, 2 - Sauls et al., 2019, 3 - Jun lab unpublished results. See Materials and Methods for our analysis methods.

### The correlation structure in the MG1655 strain in Si & Le Treut et al. (2019) is due to pyrimidine pseudo-auxotrophy

While six out of nine experiments were in favor of the division-centric model, there were still three experiments that were more consistent with the initiation-centric model (which includes the aforementioned experiment in which the adder phenotype breaks; Figure 3). However, we noticed that the three experiments used the same strain with the genetic background of *E. coli* K-12 MG1655. Kustu and colleagues extensively studied the growth physiology of MG1655 (Soupene et al. 2003). Among the several growth defects, *rph-1* is known to cause pyrimidine pseudo-auxotrophy in MG1655. While pyrimidine pseudo-auxotrophy *per se* does not affect the adder phenotype as shown before (Si and Le Treut et al. 2019; Campos et al. 2014), it could affect correlation structures involving other physiological parameters. Fortunately, in MG1655, pyrimidine pseudo-auxotrophy can be alleviated by supplementing the growth media with uracil (Soupene et al. 2003). Indeed, the three experiments supporting the initiation-centric model lacked the uracil supplement.

Following the study by Kustu and colleagues, we repeated our experiments using uracil supplemented media. This reverted the correlation structure of the three experiments, and the MG1655 also supported the division-centric model (Figure 3). These results suggest that pyrimidine pseudo-auxotrophy affects the correlation structure observed in MG1655 in Si & Le Treut et al. (2019). With the physiological defect fixed, all our experiments in both *E. coli* and *B. subtilis* unanimously supported the division-centric model.

### The *E. coli* BW strains also support the division-centric model

We next asked why Witz *et al*.’s own data supported their initiation-centric model, and repeated the ***I***-value analysis of all their pre-processed four datasets (Materials and Methods). To our surprise, our own analysis showed that their data are in fact more consistent with the division-centric mode (Figure 4A).

**Figure 4:**
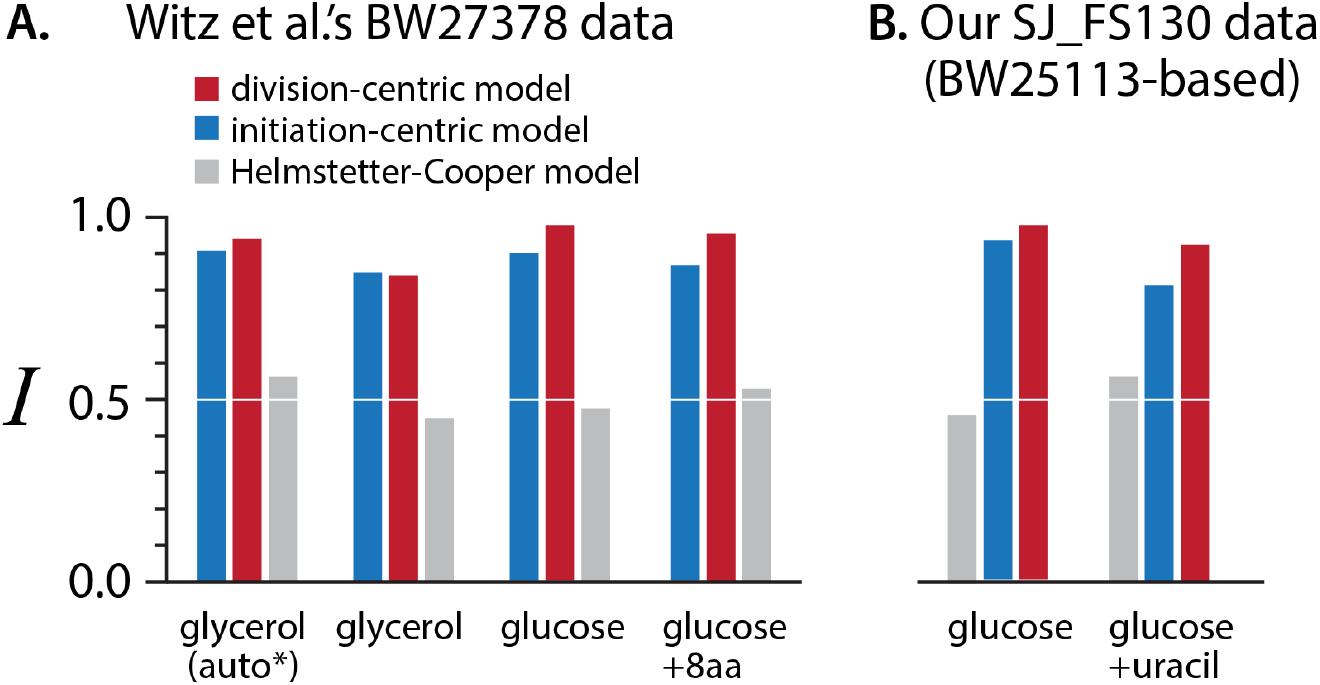
Results of the BW strains used in Witz *et al*. (**A**) and our new experiment (**B**). The glycerol (auto*) is the same experiment as glycerol, but Witz et al. used automated image analysis for detecting replication initiation.

To ensure our results are correct, we also repeated experiments using an *E. coli* K-12 strain with a genetic background practically identical to the one used in Witz et al. Two notable differences in experimental methods between our two studies are the choice of strains and how replication cycles were tracked at the single-cell level. For example, Witz *et al*. used *E. coli* BW27378. This is one of the widely adopted strains because it was used by Barry Wanner (BW) for the development of the Λ red system (Datsenko and Wanner 2000) and subsequently the Keio collection (Baba et al. 2006). Incidentally, BW27378 also has the rph-1 mutation, although it is not known whether the strain also exhibits pyrimidine pseudo-auxotrophy as in MG1655.

To track replication cycles, Witz *et al*. used FROS (fluorescent repressor-operator system) with a well-known LacI-mVenus and the LacO cassette inserted near *oriC*. This method in principle allows detection of replication initiation by measuring the timing the duplicated origin splits. There are two main caveats with this approach: (1) the measured initiation timing is almost always delayed by an uncontrolled amount of time because of the extensively studied phenomena of origin cohesion (Lesterlin et al. 2012; Sunako, Onogi, and Hiraga 2001), i.e., duplicated origins stay together for different durations of time under different growth conditions (2) this method cannot detect termination of replication. By contrast, the replisome-based tracking of DNA replication does not suffer from these caveats.

We thus transduced *dnaN-YPet* (encodes for the β sliding clamp in the replisome) to BW25113 (almost identical genetic background as Witz *et al*.’s BW27378) and repeated the experiments (Materials and Methods). Consistent with our analysis results of Witz *et al*.’s data, we found that data from these new experiments also supported the division-centric model (Figure 4B), with or without uracil supplement. While it is beyond the scope of this work to study details of physiological properties specific to the BW strains, it is clear that all experimental results support the division-centric model.

In the following two sections, we switch our focus from experiment to modeling, and show that the initiation-centric model based on the four parameters derived in Witz et al. (2019) cannot reproduce size homeostasis by the adder principle.

### The I-value analysis is insensitive to differences between size-control models

While our analysis shows that the single-cell data is more consistent with the division-centric model than the initiation-centric model, the ***I*** values of both models were close to 1 and their differences were typically less than 10%. This raises the question whether the ***I***-value analysis can really identify the correct size control model based on the ***I*** values. Therefore, we have computed the ***I*** values for all combinations of 3 among 18 variables (total of 816) using the experimental dataset analyzed by Witz and colleagues (their figure 7). The division-centric and the replication-centric models had high scores, ranking 57th and 88th respectively, yet other combinations of variables had even higher ***I*** values. In addition to the one condition analyzed by Witz et al, we also performed the analysis on the other 3 conditions of their study. Overall, we found that the division-centric model always had a higher ***I*** value than the replication centric model with only one exception of glycerol condition (Figure 4A). We also repeated the analysis using experimental data from the 6 conditions published in Si & Le Treut (2019), and obtained similar results.

Most importantly, the rank-ordered ***I*** values showed smooth changes from the highest value to the lowest value from the data in both studies. For this type of dimensional reduction approach to be useful (such as the principal component analysis), the ***I*** values ideally should show an abrupt transition such that only a small fraction of the models have significantly higher ***I*** values than the rest. Based on our results, we conclude that the ***I***-value analysis cannot reveal true (independent) control parameters.

**Figure 5:**
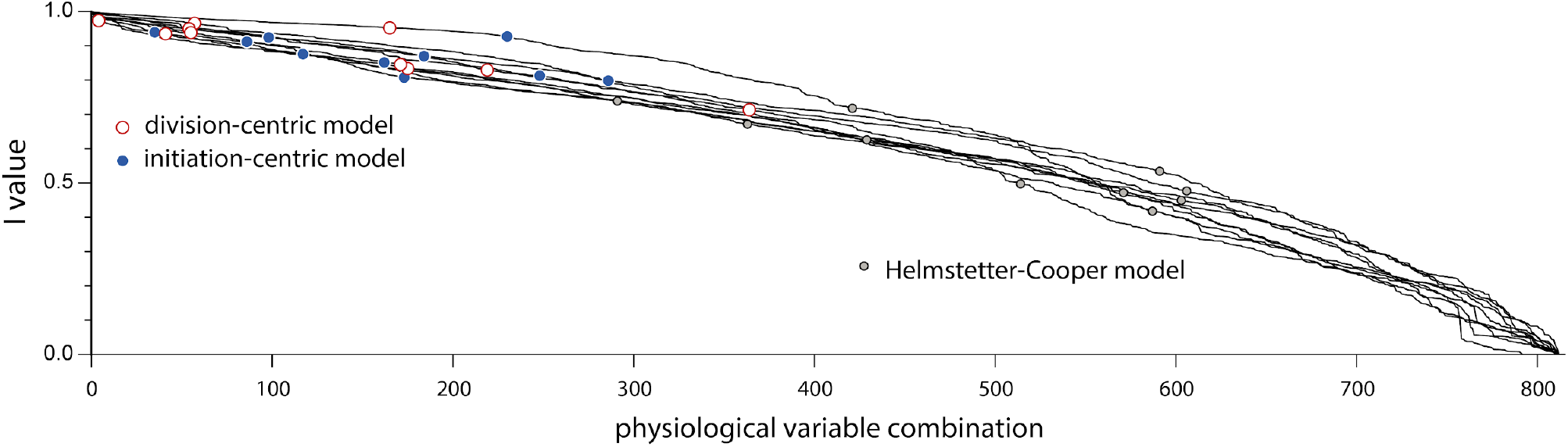
Ranked-order ***I*** values for combinations of 3 physiological variables for a total of 10 single-cell experiments: 4 experiments from Witz et *al* (2019) and for 6 experiments from Si & Le Treut et *al* (2019).

### The replication-centric model by Witz *et al*. is incompatible with size homeostasis by adder

In the presence of physiological fluctuations, the cell-size homeostasis mechanism ensures the return of cell-size to its steady-state value throughout several cell generations. A crucial quantity in cell size homeostasis is the mother/daughter (Pearson) correlation, ρ_d_, for cell size at division. The adder principle is equivalent to having a correlation coefficient ρ_d_ = 1/2 as shown by many (Amir 2014; Taheri-Araghi et al. 2015; Voorn, Koppes, and Grover 1993; Material and Methods). However, the initiation-centric model by Witz *et al*. does not produce this result. Specifically, we obtained

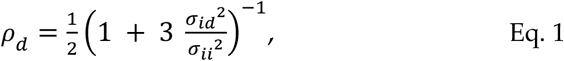

where σ_id_ and σ_ii_ are the standard deviations of δ_id_ and δ_ii_, respectively (see Materials and Methods for derivation). Therefore, this correlation coefficient is always smaller than 1/2, and it only reproduces the “adder” behavior in the deterministic limit σ_id_ → 0. The range of experimental values of σ_id_/σ_ii_ is 0.5-1.3 in Witz *et al*.’s study and 0.6-2.7 in Si & Le Treut (2019). Therefore, the model by Witz et al. predicts significant deviations from the adder phenotype observed in experiments [data summarized in (Jun et al. 2018)], much closer to a sizer model (ρ_d_ = 0; Figure 6). We further confirmed these predictions using simulations (see the Simulation section in Materials and Methods).

**Figure 6:**
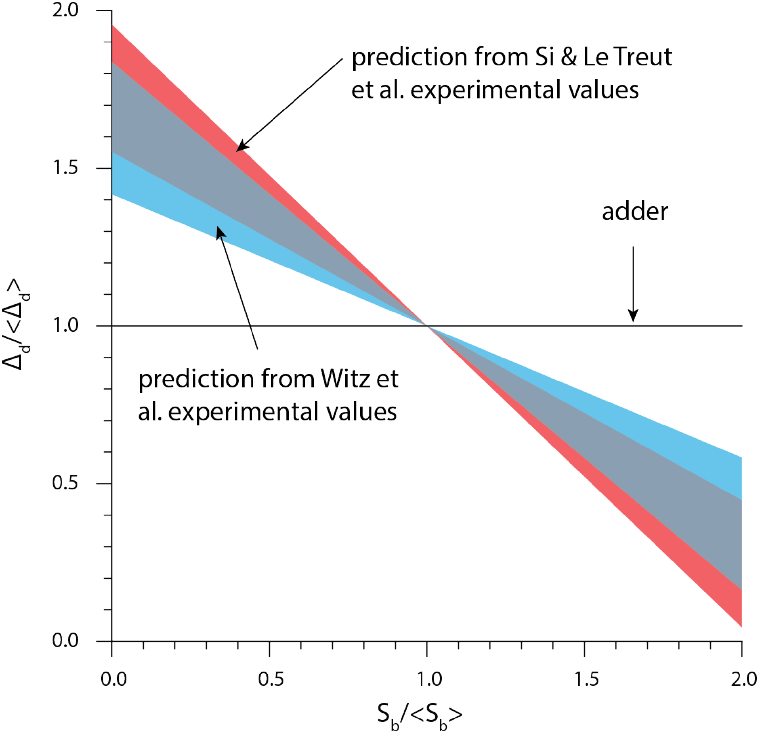
The theory based on the initiation-centric model predicts a more sizer-like behavior. We used Eq. 1 and the experimental values of σ_id_/σ_ii_, from Witz et al. (2019) and Si & Le Treut et al. (2019).

## Conclusion and outlook

Correlation analysis has become a powerful tool to interpret the flood of biological data enabled with the advent of high-throughput measurements. Correlations can indeed be a good starting point for a mechanistic model of the biological process under investigation. A case in point is the discovery of the original adder principle: the systematic lack of correlation between birth size and size added from birth to division in many conditions led to the formulation of a phenomenological model explaining how cell size converges toward a steady-state value.

However, correlation analysis can involve many variables, and the resulting complexity can be daunting and elude the original biological question. For example, in this work we showed that the model formulated on the basis of a correlation analysis does not in fact reproduce the adder principle. It is also a challenge to assess how sensitive on the experimental conditions the correlations are.

For this reason, a good strategy is to focus on correlations that are more robust than others across several conditions. Finding the control parameters that can change a seemingly invariant correlation should indeed give serious clues about the molecular mechanism that is responsible for it. Ideally, the control parameters should be linked to mechanistic hypotheses, such that they should change in response to biological perturbations that can be predicted and tested experimentally. We previously presented one such example to explain the mechanisms of cell-size control in bacteria (Si and Le Treut et al. 2019). We envision many other examples to emerge in connecting phenomenology and mechanisms in quantitative microbial physiology and beyond.

## Materials and Methods

### Availability of analysis, results, and note

All the numerical results, including simulations, discussed in the main text are available with detailed notes in the GitHub repository: https://github.com/junlabucsd/DoubleAdderArticle.

### Strains and growth conditions

The following strains were used in this study.

- BW25113: F-DE(araD-araB)567 lacZ4787(del)::rrnB-3 LAM-rph-1 DE(rhaD-rhaB)568 hsdR514. [Note that the genotype of BW27378 is F-DE(araD-araB)567 lacZ4787(del)::rrnB-3 LAM-rph-1 DE(rhaD-rhaB)568 hsdR514 DE(araH-araF)570(::FRT).]
- SJ_FS130: To construct this strain, we introduced ΔdnaN::[dnaN-ypet] into BW25113 using P1 transduction.
- SJ_DL188: MG1655 F-λ-rph-1 dnaA msfGFP kan mCherry-dnaN

We used minimal MOPS media for the MG1655 experiments in Figure 3 and minimal M9 glucose media with and without uracil for the SJ_FS130 (BW25113 based) experiments in Figure 4.

### Microfluidics, Microscopy and Image processing

We used the same method as described in Si & Le Treut et al (2019).

### *I*-value analysis

Following the methodology proposed by Witz and colleagues, we first defined the covariance matrix K = [k_ij_] of the analyzed variables. Then the score was computed as:

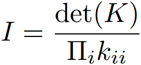

where the k_ii_ are the diagonal elements of the covariance matrix, hence the variances.

We defined 18 physiological variables in line with the definitions given by Witz *et al*. (see Figure S1): λ, S_b_, S_d_, S_i_, Δ_d_, Δ_bi_, Λ_i_, Λ_*i*^(n-1)^_, Λ_i^(n+1)^_, δ_ii_, δ_id_, R_ii_, R_id_, R_bd_, R_bi_, τ, τ_cyc_ and τ_ii_. We generated all possible combinations of 3 variables, that is 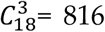. We emphasize the difference between Λ_i_ and S_i_. The former is the cell size per origin of replication at the initiation event associated with the division in the current generation; therefore it could be in a previous generation (*e.g*. mother, grand-mother cell). The latter is the cell size when replication initiation occurs in the current generation. These 2 variables are only the same in the case of non-overlapping cell cycles.

We also found that the results of this analysis were sensitive to processing of the experimental data. In their work, Witz and colleagues, instead of using the measured values for S_b_, S_d_, S_i_ and Λ_i_, first fitted traces of cell sizes to an exponential function and took values interpolated by this fit. Yet it did not affect the relative ranking of the division-centric model with respect to the initiation-centric model.

#### Parameters in the initiation-centric model by Witz et al

In their analysis, Witz *et al*. used 4 variables: Λ_i_, δ_ii_, δ_id_ and λ. However, Λ_i_ is not strictly required. Indeed, from the knowledge of Λ_i_ in the first cell of the lineage, one may reconstruct the subsequent replication and division cycles from the sole knowledge of δ_ii_, δ_id_ and λ. Therefore Λ_i_ appears to be an initial condition, and the other 3 variables actually define a model for division and replication cycle progression.

#### Parameters in our division-centric model and in the Helmstetter-Cooper model

In our division-centric model, the cell size at birth S_b_ is only required in the first cell of the lineage. The knowledge of only the 3 variables δ_ii_, Δ_d_ and λ in each new generation is sufficient to reconstruct the whole lineage. The Helmstetter-Cooper model features the cell size at replication initiation per origin s_i_, the duration of the cell cycle τ_cyc_ and the growth rate Λ. Similarly to our model, the cell size at birth Sb is only required in the first cell of the lineage. Then, from this initial condition and the knowledge of the 3 variables s_i_, τ_cyc_ and Λ at each new generation, one can reconstruct the whole cell lineage.

In summary, only 3 variables are necessary and sufficient to self-consistently define the cell cycle and cell size. This is why we carried out this “determinant” analysis using each of the 3-variable sets that we just described. Nevertheless, we also examined inclusion of a fourth variable, and found that it did not affect our conclusions.

### Theory: cell size homeostasis in Witz *et al*.’s initiation-centric model

In the model proposed by Witz and colleagues, the cell size per origin at division is determined by the cell size at initiation per origin Λ_i_, the added size per origin between consecutive replication initiation events δ_ii_, and the added size per origin from replication initiation to cell division δ_id_. The following relation holds:

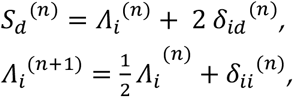

where the index as *n* denotes the generation (or division cycle). Under the assumption that δ_ii_^(n)^ (resp. δ_id_^(n)^) are independently and identically distributed Gaussian stochastic variables with mean μ_ii_ (resp. μ_id_) and standard deviation σ_ii_ (resp. σ_id_), it follows that Λ_i_^(n)^ and S_d_^(n)^ are also Gaussian stochastic variables. At large *n*, they converge to the limiting distributions Λ_i_ ≡ N(μ_i_, σ_i_) and Sd ≡ N(μ_d_, σ_d_), where:

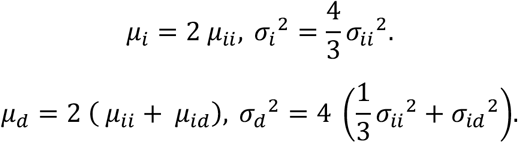

The mother/daughter correlation for division size is a central quantity in cell size homeostasis, which can be derived in this model. As a first step, let us define the centered variables: dΛ_i_^(n)^ = Λ_i_^(n)^ - μ_i_ and dS_d_^(n)^ = S_d_^(n)^ - μ_d_. We then obtain the relations:

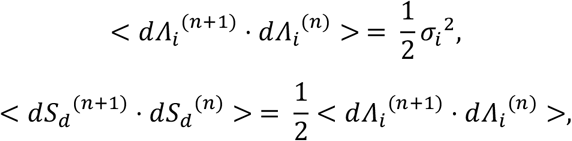

where the brackets denote averages. We therefore obtain the mother/daughter Pearson correlation coefficients (in the large *n* limit):

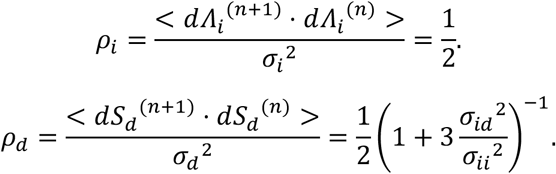

In this model the joint distribution (S_d_^(n)^, S_d_^(n-1)^) is a bivariate Gaussian, therefore we can write the conditional expectation of S_d_^(n)^ as:

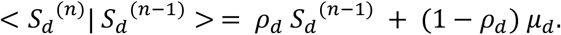

With the hypothesis of symmetrical division, namely S_b_^(n)^ = 2 S_d_^(n-1)^, we obtain for the conditional expectation of the added size from birth to division:

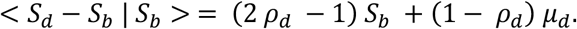

Therefore, the “adder” principle is equivalent to having *ρ*_d_ = 1/2. From these results, the model proposed by Witz and colleagues always results in 2 *ρ*_d_ - 1 < 0. In fact, it only reproduces the “adder” principle in the deterministic limit σ_id_ → 0.

### Simulations

In this study we have performed simulations of both initiation-centric and division-centric models. For this, we have re-used the code provided by Witz and colleagues (2019). Few and minor modifications have been made, but these modifications did not affect the outcome or the essence of the original simulations. Simulations performed consist of:

- Repeats of the simulations performed by Witz and colleagues in their original study.
- Simulations using Witz and colleagues original parameters, but with perfectly symmetrical divisioni.
- Simulations of Witz et al. model, and our model, using experimental parameters taken from experimental datasets from Witz and et *al*. and datasets published in (Si and Le Treut et al. 2019).

See https://github.com/junlabucsd/DoubleAdderArticle for more details.

#### Deviations of Witz *et al*.’s initiation-centric model (simulation) from the adder behavior

Witz *et al*.’s simulation reproduced the adder behavior observed in their data, in apparent contradiction with our prediction in Eq. 1 that their initiation-centric model is inconsistent with size homeostasis by the adder. We analyzed their simulations and found that they produced the adder-like behavior because of the additional fluctuations in the septum position (Figure S2A).

Experimentally, septum position represents the most precise control among all measured single-cell parameters with CV < 5% (Taheri-Araghi et al. 2015; Sauls et al. 2019). Indeed, removing fluctuations in the septum position alone made the simulation deviated from experiment in a quantitative manner consistent with Eq. 1. We also conducted a similar analysis using our experimental data (Si and Le Treut et al. 2019), and reached the same conclusion (Figure S2B). Based on this observation, we conclude that the model proposed by Witz *et al*. does not self-consistently explain the adder phenotype.

---

**Figure S1:**
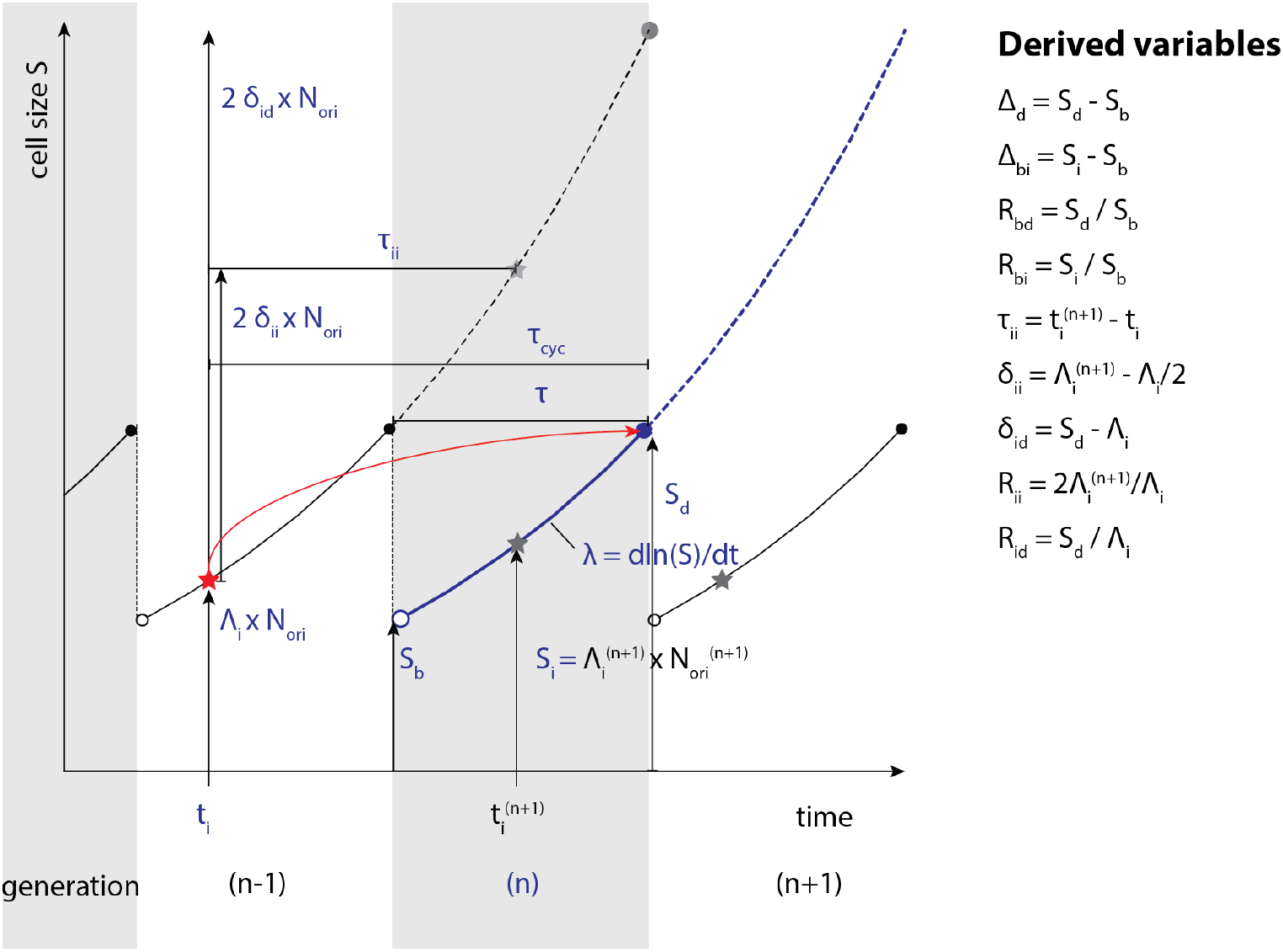
Definition of the physiological variables in a scenario with 2 overlapping cell cycles. All variables with blue font are associated with the current generation (*n*). We added a superscript when using physiological variables associated with another generation. We also defined variables derived from these quantities. Replication initiations are indicated with stars, and the red star is the initiation determining the division for the current generation. N_ori_ is the number of origins of replication just before initiation happens.

**Figure S2:**
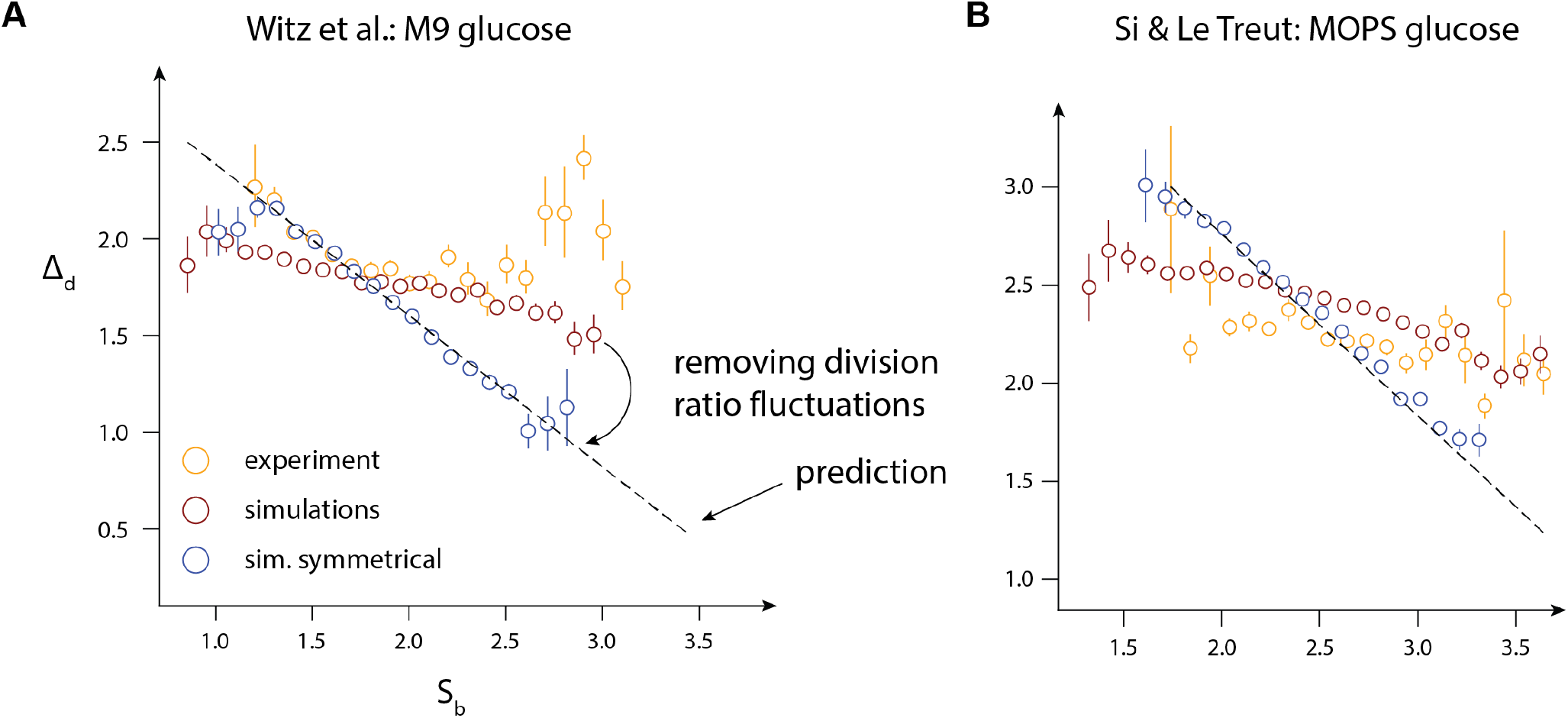
Agreement of the initiation-centric model with experimental data from **A.** Witz et al., M9 + glucose condition and **B.** Si & Le Treut et al., MOPS + glucose condition. **A.** The agreement of simulations with experimental data gets worse after removing fluctuations in the division ratio. **B.** The agreement of simulations with experimental data is not as good, and gets worse after removing fluctuations in the division ratio.

